# PhenoBR: a model to phenotype body condition dynamics in meat sheep

**DOI:** 10.1101/2020.12.01.407098

**Authors:** Tiphaine Macé, Eliel González-García, György Kövér, Dominique Hazard, Masoomeh Taghipoor

## Abstract

In situations of negative energy balance (NEB) due to feed scarcity or high physiological demands, body energy reserves (BR), mainly stored in adipose tissues, become the main sources of energy for ruminants. The capacity to mobilize and restore such BRs in response to different challenges is of major concern in the current context of breeding for resilience. Body condition score (BCS) is a common, practical indicator of BR variations throughout successive productive cycles, and quantitative tools for characterizing such dynamics at the individual level are still lacking. The main objective of this work was to characterize body condition dynamics in terms of BR mobilization and accretion capacities of meat sheep during their productive lifespan through a modelling approach.

The animal model used in this work was the reproductive meat ewe (*n* = 1478) reared in extensive rangeland. Regular measurements of BCS for each productive cycle were used as the indicator of BR variations. A hybrid mathematical model and a web interface, called PhenoBR, was developed to characterize ewes’ BCS variations through four synthetic and biologically meaningful parameters for each productive cycle *i*: BR accretion rate 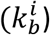, BR mobilization rate 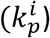, plus the time of onset and the duration of the BR mobilization, 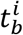 and Δ*T^i^*, respectively.

The model converged for all the ewes included in the analysis. Estimation of the parameters indicated the inter-individual variability for BR accretion and mobilization rates, and for the length of the mobilization period. Body reserve mobilization rates were closely correlated between productive cycles. Significant correlations between BR mobilization and accretion rates suggest that the two processes are biologically linked. Parameters *k_p_* and *k_b_* decreased as parity increased. BR mobilization rate and duration increased as litter size increased, while BR accretion rate decreased.

Individual characterization of animals by these parameters makes it possible to rank them for their efficiency in the use of body reserves when facing NEB challenges. Such parameters could contribute to better management and decision-making by farmers and advisors, e.g. by adapting feeding systems to the individual characteristics of BR dynamics, or by geneticists as criteria to develop future animal breeding programs including BR dynamics for more robust and resilient animals.

## Introduction

Body energy reserves (BR) are the main source of energy in ruminants facing negative energy balance (NEB) challenges such as highly demanding reproductive cycles or feed scarcity periods (1,2). The capacity of ruminants to mobilize and restore such BRs in response to challenges is of major concern in the current context of breeding for robustness and resilience, and for the ultimate sustainability goals of farming systems. Priorities chosen by complex mechanisms related to nutrient allocation and trade-offs become critical at the individual level. BR administration and operational feeding strategies are among the main resources set by the animal and the farmer, respectively, to cope with such challenges.

In meat sheep, a broad intra-flock variability in the dynamic of BR was shown during their lifespan including several reproductive cycles. The genetic determinism related to the BR changes has been demonstrated by considering successive physiological stages independently (3,4). The BR dynamics could be considered in selection strategies designed to improve the ewes’ adaptive capacities while maintaining production and welfare. It is therefore of major interest to phenotype ewes for their capacity of resilience in situations of NEB.

Despite the importance of BR dynamics for animal robustness, its use in animal husbandry and breeding is still at its very early stages (5,6). There are two main reasons for this. Firstly, there is a lack of dynamic data describing animal response facing different, successive challenges of different magnitude and amplitude. Progress in technology in the last decades has made it possible to measure several indicators of animal performance with relatively high frequency (i.e. dynamic records). Through the development of new monitoring technologies, dynamic measures of indicators such as body weight (BW), milk yield and feed intake are available with higher frequency and at lower cost. Secondly, the absence of synthetic criteria for characterizing and quantifying BR dynamics is still a limiting factor for the use of such a complex phenotype. Body condition score (BCS) is a practical and conventionally used variable for monitoring BR dynamics. Compared to other variables such as BW, BCS is a proven reliable indicator of an animal’s fat reserves (7,8). Although some automatic advances are available, BCS continues to be a subjective but effective variable, measured in general through direct observations carried out by a trained observer (9,10). In outdoor farming systems (e.g. grazing, rangelands, etc.) there are still limitations to overcome regarding, for example, the availability of sound tools for implementing automatic measurement of BCS at the individual level.

Macé and collaborators concluded on the usefulness of BCS and BW data to study the capacity of BR mobilization and accretion in meat ewes (3). Based on mean trajectories of BCS and BW, they defined periods of mobilization and accretion throughout ewes’ lifespans and studied variations of BCS and BW in different physiological stages during these periods. Some dynamic and mechanistic models were developed to predict fat fluxes in cattle during transition periods, using BCS and DMI (11) and to describe BR variations in dairy goats during their lifespan using daily measurement of BW (12). However, to our knowledge, there is no tool for individual treatment of BCS data in sheep to automatically detect periods of mobilization and accretion and to quantify individual response to NEB challenges, in terms of BR mobilization and accretion rates. Moreover, existing longitudinal modelling in genetics is ill-adapted to considering dynamic processes of BR, especially in the case of low frequency phenotyping.

The objective of the present work was to produce a quantitative, integrated tool for characterizing the BR dynamics of meat ewes reared outdoors with a small number of synthetic variables. To this end, a dynamic model, called PhenoBR, was developed, based on a system of ordinary differential equations, as a support to describe BCS variations in meat ewes over several productive cycles. This model is implemented in a web interface (http://adaptive-capacity.herokuapp.com/) that lets users (even those unfamiliar with modelling) test different hypotheses and results of the model. This tool should be very useful for investigating physiological and genetic components of BR dynamics and developing future animal breeding programs for more robust and resilient animals.

## Materials and Methods

The animal model used in this work was the reproductive meat ewe, during its whole lifespan. However, the model can be considered as generic and adaptable to other ruminant species and productive purposes. The parameters of this model represent the ewe’s characteristics in terms of its capacities to mobilize and restore BR at each production cycle. Two main types of parameters are expected. The first category comprises time-related parameters (i.e. time related to the beginning of BR mobilization and the interval, or duration of the BR mobilization period for each productive cycle). The second category of parameters relate to the intensity of the BR mobilization period and the capacity to recover an expected BR status. The model is hybrid in that it combines a data analysis procedure and a more concept-driven model.

### Ethics statement

This study was conducted without carrying out any additional animal experiments or biological sampling. The phenotypes used in the present study had been collected previously in other projects that fully complied with applicable legislation on research on animals in accordance with the European Union Council directive (2010/63/EU). All the experimental procedures were performed according to the guidelines for the care and use of experimental animals and were approved by the local ethics committee (approval number APAFIS4597-2016031819254696).

### Animals and experimental data

The datasets used in the present study have been previously described in detail (3,4). Briefly, the experimental animals belonged to the Romane meat sheep breed. The monitored ewes *(n* = 1478) were reared under extensive conditions on the rangelands of the INRAE La Fage experimental farm (Causse du Larzac 43°54’54.52”N; 3°05’38.11”E, Roquefort du Soulzon, France). Ewes performed one productive cycle per year. The biological productive cycle length was thus 365 days. Data were collected for the period 2002 to 2015.

In this study, the BCS was used as the main indicator to illustrate the BR dynamics (i.e. the capacity of ewes to mobilize and restore BRs). Ewes were measured for BCS, with eight measurements collected regularly during each female’s productive cycle according to a physiological stage schedule. Ewes were measured during one to three entire productive cycles (1278 ewes for cycle 1, 1204 for cycle 2 and 521 for cycle 3). All the ewes included in the present study were pregnant and suckled at least one lamb until weaning (~80 days after lambing). To assess BCS, the original grid described by Russel and coworkers (7) was used and subdivided into a 1/10 scale, i.e. from 1 to 5 with 0.1 increments. All the measurements were recorded in the Geedoc database (https://germinal.toulouse.inra.fr/~mcbatut/GEEDOC/).

### Modelling procedure: General model

To characterize the ewes’ response to the increasing energy requirements at each productive cycle in terms of BCS variations, a system of ordinary differential equations was developed. For present purposes, the significant increase in energy requirements during a productive cycle is termed perturbation (physiological challenge due to pregnancy and suckling). The objective of model development is to describe the ewe’s response to that perturbation using BCS variation as the indicator of ewe’s response. Two interrelated state variables were used to describe the ewe’s response at productive cycle *i*: *p_i_* for the decrease in BCS during the mobilization period and *BCS_i_* for the variations in BCS during the same productive cycle including its decrease by *p_i_* and its recovery (Figure 1). The driving force of this model is the intrinsic capacity of the ewes to maintain or to restore their BRs to reach the expected BCS, considered as the value of BCS in the absence of perturbations, noted *BCS_m_*. The concept of the expected trajectory of performance is already described in the literature (13). In the database used, the ewe population attained at most three productive cycles, and the total BCS variations was calculated as the sum of *BCS_i_* variations, for *i* ∈ {1,2,3}.

**Figure 1.**
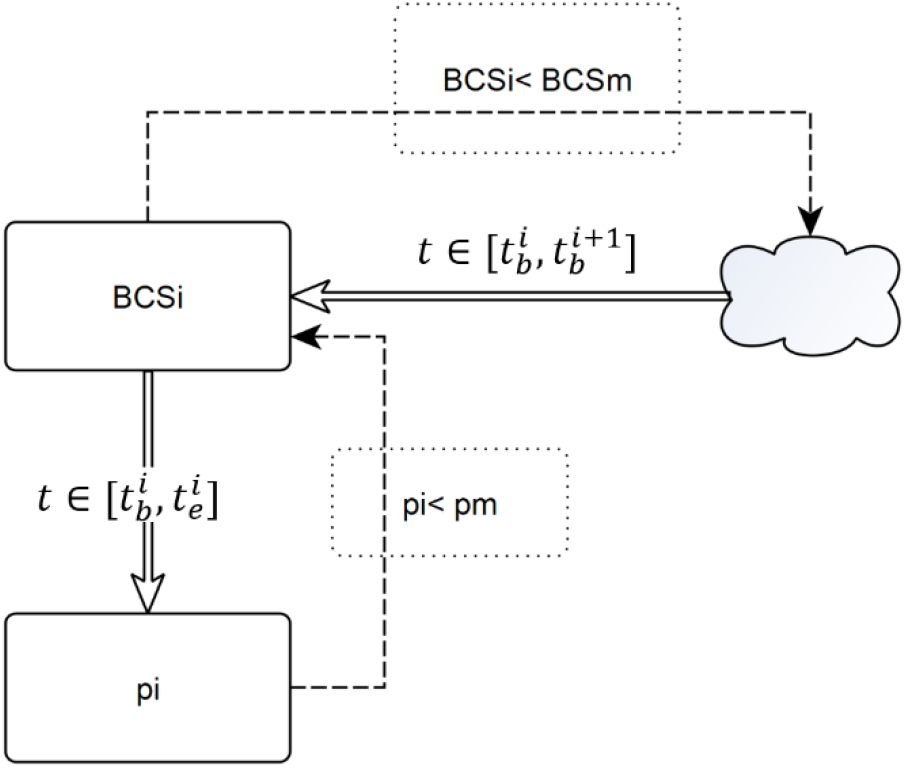
The model for one productive cycle. Flux to BCS is regulated by the difference between *BCS_i_* and the *BCS_m_* for a given animal. The flux to *p_i_* is activated in the interval 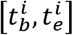 and will stop when it reaches *p_m_*. From the beginning of the perturbation, the decrease in *BCS_i_* is counterbalanced by all internal physiological mechanisms of the ewes seeking to keep the *BCS_i_* close to *BCS_m_*.

The general model for one productive cycle could then be written as Equation 1.

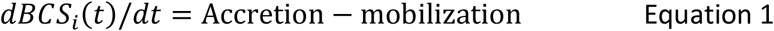

For productive cycle *i*, periods of BR mobilization and accretion are both assumed to start from the point when *BCS_i_* starts to decrease 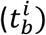, considered as the starting point of the perturbation being studied. In this study, it is assumed that BCS is a good indicator of the ewes’ BR variations, as previously demonstrated. When the animal is no longer able to meet the energy demands induced by pregnancy and lactation using only the energy available from ingested feeds, it will mobilize the energy stored in its adipose tissues (i.e. its energetic BRs). However, during and after this period, the animal uses internal mechanisms to limit or to compensate for the quantity of BRs used during this period. This is illustrated by the effort of the animal to reach *BCS_m_*. To clearly separate the effects of each productive cycle, the BR recovery capacity associated with productive cycle *i* starts at 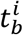 and lasts until the beginning of the next productive cycle 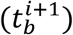.

The BR mobilization period is assumed to be over at 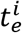, which is the end of the BCS decrease period (i.e. end of perturbation*i*). 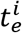 is associated with the point where the capacity of BR accretion surpasses the ewe’s energy requirement. Equations 2.a and 2.b describe the variations in *BCS_i_* and the simultaneity and continuity of BR mobilization and accretion processes for productive cycle *i*.

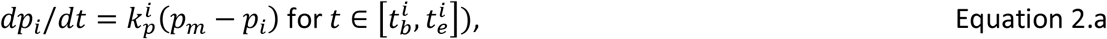

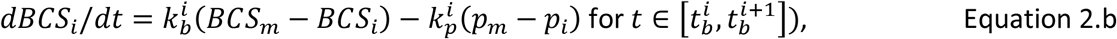

Where 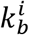 and 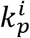 are the rates for BR accretion and mobilization during the productive cycle *i*, respectively. The constant *p_m_* stands for the maximal loss of BCS due to the perturbation. Variations in total BCS during the ewe’s lifespan are the sum of variations of *BCS_i_* at each productive cycle as stated in Equation 3.

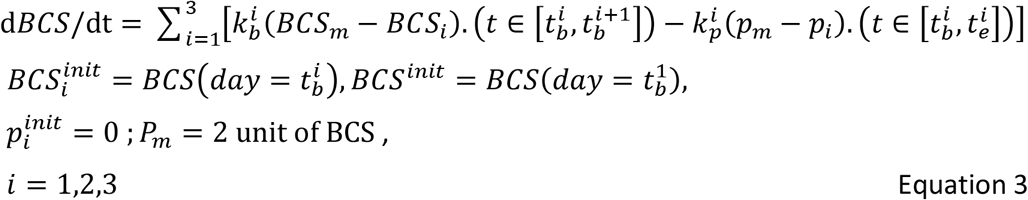

where 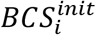 is the initial value of *BCS_i_*, and *BCS^init^* is the initial value of the total *BCS*. Using this model, the response of the animal at each productive cycle is characterized by the set of parameters 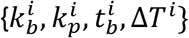, where 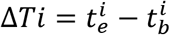, and stands for the duration of the mobilization period. Parameters 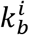 and 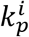 are estimated using the minimization function in Equation 4

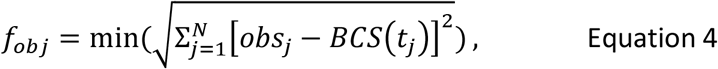

where *j* stands for the number of observation points *obs_j_* of BCS during the ewe’s lifespan, and *BCS*(*t_j_*) is the estimation of BCS at age *t_j_* by the model (Equation 3). *N* is the number of observations for a given animal (at most *N* = 24).

Other parameters of the model are time-related 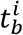 and Δ*T^i^*, which are determined using a data analysis procedure for automatic detection of the perturbation within each productive cycle. The dataset involved ewes with 1, 2 or 3 parities, and the data analysis procedure allowed automatic determination of the number of parities to be considered for each ewe, and the beginning and the duration of the associated perturbation.

### Modelling procedure: Automatic detection of perturbation period

The beginning and end of each productive cycle was taken to be from one mating to the subsequent one. When all data were missing in a given period, it was taken as a missing productive cycle. In this section, a data analysis procedure based on functional data analysis (FDA) was developed to automate detection of perturbations within each productive cycle. In this procedure, the beginning of a perturbation 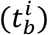 is where the observed BCS started to deviate from the initial value of *BCS_i_* 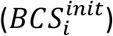. The end of the perturbation 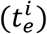 is where it started to recover.

The expected value of BCS (*BCS_m_*) is determined depending on ewes’ maximum value of BCS *(BCS_max_*). If *BCS_max_* < 3.4 then *BCS_m_* = 3.5, otherwise *BCS_m_* = 4. The value of 3.4 was chosen because it was the common observed maximum BCS value for most of the ewes in the dataset. Also, in the construction of PhenoBR, the value of threshold *BCS_m_* will be approached but never reached. It should therefore be larger than *BCS_max_*, to let ewes reach their *BCS_max_*. A B-Spline regression with smoothing parameters *λ* = 10^3^ was fitted to the BCS data of each animal. This regression is drawn from methods on analysis of functional data (13–15), in which the flexibility of the fitted function is ensured by the use of the piecewise polynomial (in this case of degree 3) and the smoothness is adjusted by the definition of a roughness penalty *λ*. This regression is in particular useful in cases where the general shape of the function under study is unknown, which is the case in the presence of perturbations. Equation 5 describes the objective function to fit the B-spline function to observed data, for the estimation of the function *BCS*(*t*).

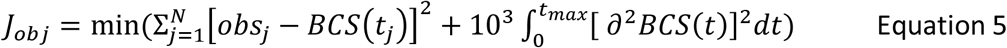

Where *obS_j_* stands for the *j*th observation of BCS, *BCS*(*t_j_*) is the estimation of BCS by the B-spline function and *t_max_* is the last day of the BCS record for each individual. Zeros of the derivative of the estimated function *BCS*(*t*) are maximums and minimums of *BCS*(*t*). Maximum indicates the beginning of potential deviations from 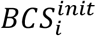 and minimums are the beginning of the recovery period (Figure 2). A limited number of exceptions should be considered if several extremums are present inside a productive cycle, or if there are no extremums, (details of these cases are provided in Supplementary Hypothesis 1).

**Figure 2.**
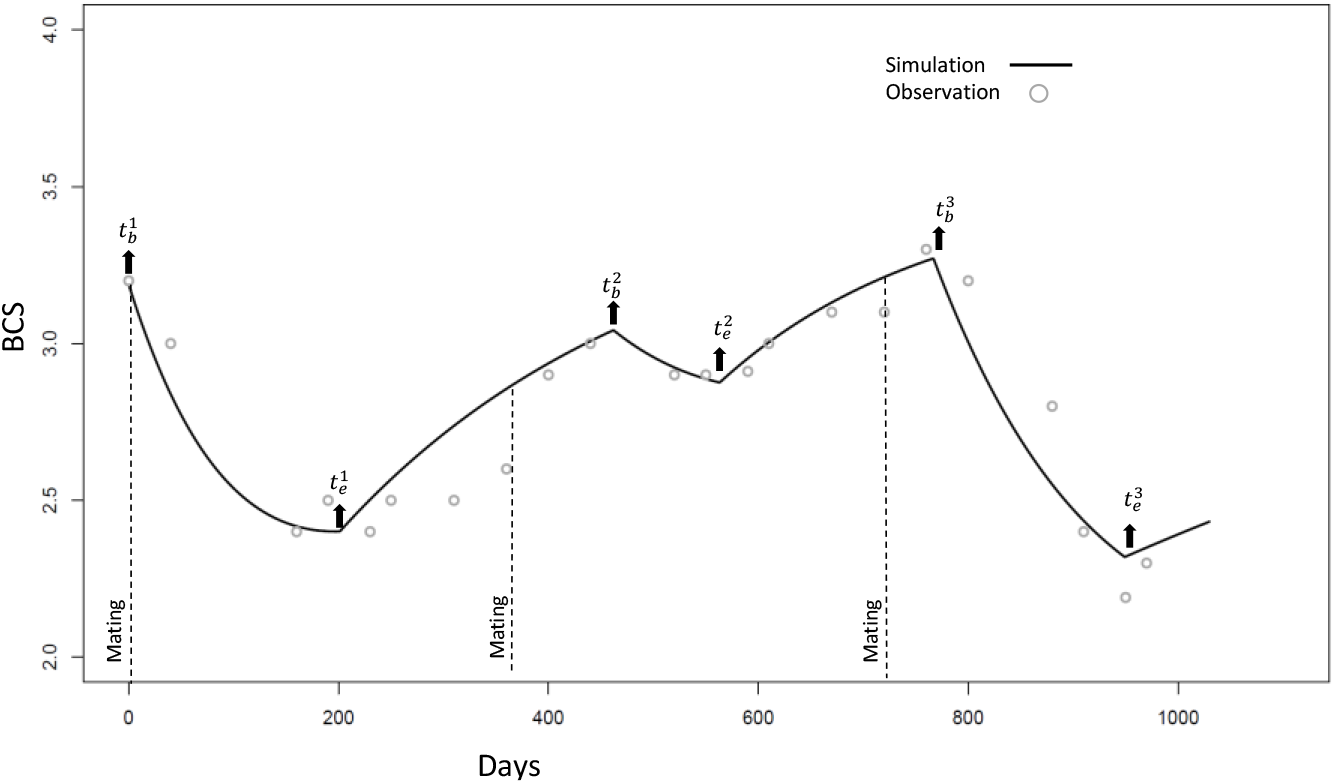
Illustration, for one ewe, of the perturbation periods as determined by the data analysis procedure. 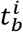 and 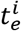 are associated with the times of the beginning and end of each perturbation. Dashed lines show the mating days of each parity as written in the original data set. Time zero represents the first BCS measurement.

Although the procedure determined the beginning and end of the mobilization period *t_b_* and *t_e_*, respectively, existing information on ewes’ physiology were also considered for a more reliable estimation of these values. In this respect, *t_b_* can take a value approximately between mating and lambing days, and *t_e_* can take a value approximately between lambing and post-weaning as defined in (3). In the construction of PhenoBR, the recovery period of each productive cycle *i* is over when the next productive period starts 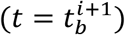. A major challenge was then to determine the recovery period when second or third productive cycles were missing for individuals from the original database. For this, it was assumed that in the absence of 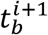, the recovery from the productive cycle *i* would be over 150 days after the postweaning period (i.e. the period corresponding to the next mating).

This procedure allowed the number of productive cycles and the associated *t_b_* and *t_e_* to be determined for each animal. All missing values of BCS were replaced by values of the function *BCS*(*t*). Figure 2 illustrates the perturbation periods as determined by the data analysis procedure against a reproductive cycle as reported in the original dataset. Table 1 summarizes all parameters of the model.

**Table 1.**
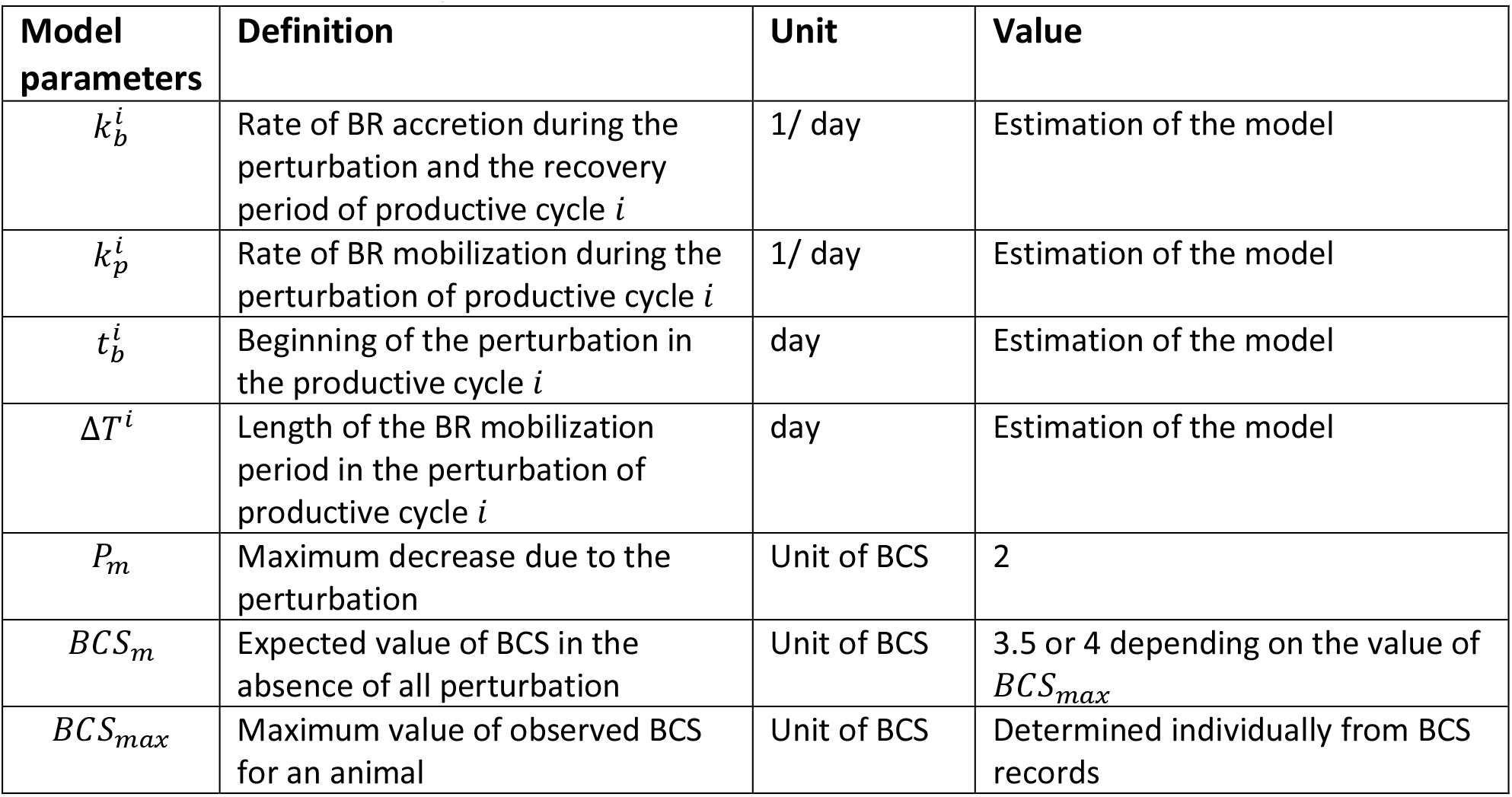
Definition of different parameters and constants used in the PhenoBR.

| Model parameters | Definition | Unit | Value |
| --- | --- | --- | --- |
| $k_b^i$ | Rate of BR accretion during the perturbation and the recovery period of productive cycle $i$ | 1/ day | Estimation of the model |
| $k_p^i$ | Rate of BR mobilization during the perturbation of productive cycle $i$ | 1/ day | Estimation of the model |
| $t_b^i$ | Beginning of the perturbation in the productive cycle $i$ | day | Estimation of the model |
| $\Delta T^i$ | Length of the BR mobilization period in the perturbation of productive cycle $i$ | day | Estimation of the model |
| $P_m$ | Maximum decrease due to the perturbation | Unit of BCS | 2 |
| $BCS_m$ | Expected value of BCS in the absence of all perturbation | Unit of BCS | 3.5 or 4 depending on the value of $BCS_{max}$ |
| $BCS_{max}$ | Maximum value of observed BCS for an animal | Unit of BCS | Determined individually from BCS records |

The software R version 3.4.2 (16) was used for the development of PhenoBR, estimation of the model parameters (function ‘optim’ of package stat) and for the data analysis procedure (package fda). The structural identifiability of dynamic model (3) for variables 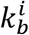 and 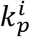 was tested using the software DAISY (17). The model was structurally identifiable, meaning that the variable estimation problem is well posed and it is theoretically possible to estimate uniquely the model variables given the available measurements (18).

### Statistical analysis

Deviations from normality were checked using the UNIVARIATE procedure of SAS (version 9.4; SAS Institute Inc., Cary, NC). None of the variables were transformed, as no major deviation from normality was observed. The Pearson correlation test was used to calculate the correlation among model variables. Analyses of variance were carried out, taking into account the repeated measures, with the MIXED procedure of SAS (version 9.4; SAS Institute Inc., Cary, NC) to test relevant effects and interactions affecting 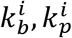 and Δ*T^i^*. Age at first lambing, parity of the ewe, litter size and year of measurement were identified as fixed effects. The age at first lambing effect is the age at which ewes lambed for the first time (i.e. 1 or 2 years; classes 1 and 2, respectively). The parity effect considered first, second and third lambing (i.e. classes 1, 2 and 3, respectively). The litter size effect took into account the number of lambs born and suckled by the ewe (i.e. singletons lambed and suckled for class 1; twins lambed but only one suckled for class 2; twins lambed and suckled for class 3 and more than two lambs lambed and suckled for class 4). The first-order interactions between productive cycle and litter size and between age at first lambing and litter size were tested. An effect was considered significant if *p* < 0.05.

Cluster analyses had been previously performed to investigate the variability of BCS profiles during each productive cycle (4). In the present study, the MIXED procedure of SAS (version 9.4; SAS Institute Inc., Cary, NC) was used to compare the parameters 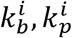 and Δ*T^i^* for the different BR clusters. For each productive cycle, models included the cluster factor together with age at first lambing, litter size and the year factors when needed.

### Software development

To make the model easily usable by non-modellers, PhenoBR is also available as an online software package on the free platform Heroku (https://www.heroku.com/). The software was developed using Python 3.7 (19) and is available on http://adaptive-capacity.herokuapp.com/

## Results

The PhenoBR fitting procedure to estimate parameters 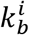 and 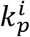 converged for the 1474 ewes in this study (*f_obj_* = 0.54±0.22 units of BCS, Equation 4). Four ewes with only one parity and less than four datapoints were removed, because of lacking enough data for estimating model parameters. The minimum and maximum of the residuals were 0.06 and 1.35 units of BCS, respectively (supplementary figure S1). The dataset used in this study contained ewes with different number of parities (from 1 to 3). To minimize the intervention of the user, PhenoBR enabled us to identify automatically the number of reproductive cycles per ewe (1, 2 or 3), and in each cycle to detect the time of the beginning 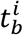, and the length of the BR mobilization periods Δ*T^i^* (Δ*T*^1^=212.3±51.0, Δ*T*^2^=174.6±53.4, Δ*T*^3^=181.1±50.2 days). Descriptive statistics of parameters estimated by the model are presented in Table 2.

**Table 2.** Descriptive statistics of parameters estimated in the modelling procedure.

| | $k_b^1$ | $k_b^2$ | $k_b^3$ | $k_p^1$ | $k_p^2$ | $k_p^3$ | $\Delta T_1$ | $\Delta T_2$ | $\Delta T_3$ | $f_{obj}$ |
| --- | --- | --- | --- | --- | --- | --- | --- | --- | --- | --- |
| Mean | 3.296 | 3.051 | 2.870 | 4.979 | 4.788 | 4.847 | 212.3 | 174.6 | 181.8 | 0.537 |
| SD | 1.034 | 1.110 | 1.078 | 1.704 | 1.661 | 1.401 | 50.9 | 53.4 | 50.2 | 0.222 |
| Min | 0.000 | 0.000 | 0.000 | 1.066 | 0.000 | 1.392 | 34.0 | 41.0 | 63.0 | 0.059 |
| Max | 7.755 | 8.548 | 8.613 | 14.349 | 16.485 | 10.494 | 341.0 | 338.0 | 372.0 | 1.355 |
| 2<sup>nd</sup> quantile | 2.458 | 2.249 | 2.257 | 3.521 | 3.428 | 3.859 | 174.2 | 135.0 | 144.0 | 0.371 |
| 4<sup>th</sup> quantile | 4.038 | 3.663 | 3.413 | 6.161 | 5.875 | 5.573 | 251.0 | 202.0 | 214.0 | 0.691 |

$k_b^i$ and $k_p^i$ are the BR accretion rate and BR mobilization rate during the productive cycle $i$ . Parameters $k_b^i$ and $k_p^i$ are all multiplied by a factor of 1000 for better clarity. Parameter $\Delta T_i = t_e^i - t_b^i$ shows the duration of the BR mobilization period at each productive cycle $i$ . The quality of model fitness is shown by $f_{obj} = \sqrt{\sum_{j=1}^N [BCS_j - y(t_j)]^2}$ which represents the residual standard error of the model.

Large positive correlations were found between parameters of BR mobilization (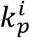 for *i* = 1, 2, 3) (Figure 3). Positive and significant *(p* < 0.005) correlations between the BR mobilization rate 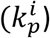 and BR accretion rate 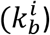 were observed for parities 1 and 2. Significant negative correlations between BR mobilization length Δ*T^i^* and rate 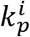 could also be observed for parities 1 and 3, i.e. the longer the BR mobilization period, the smaller the BR mobilization rate. Positive and significant correlations were also obtained between BR mobilization rates 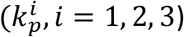. Figure 3 gives results of all ewes with different numbers of parities.

**Figure 3.**
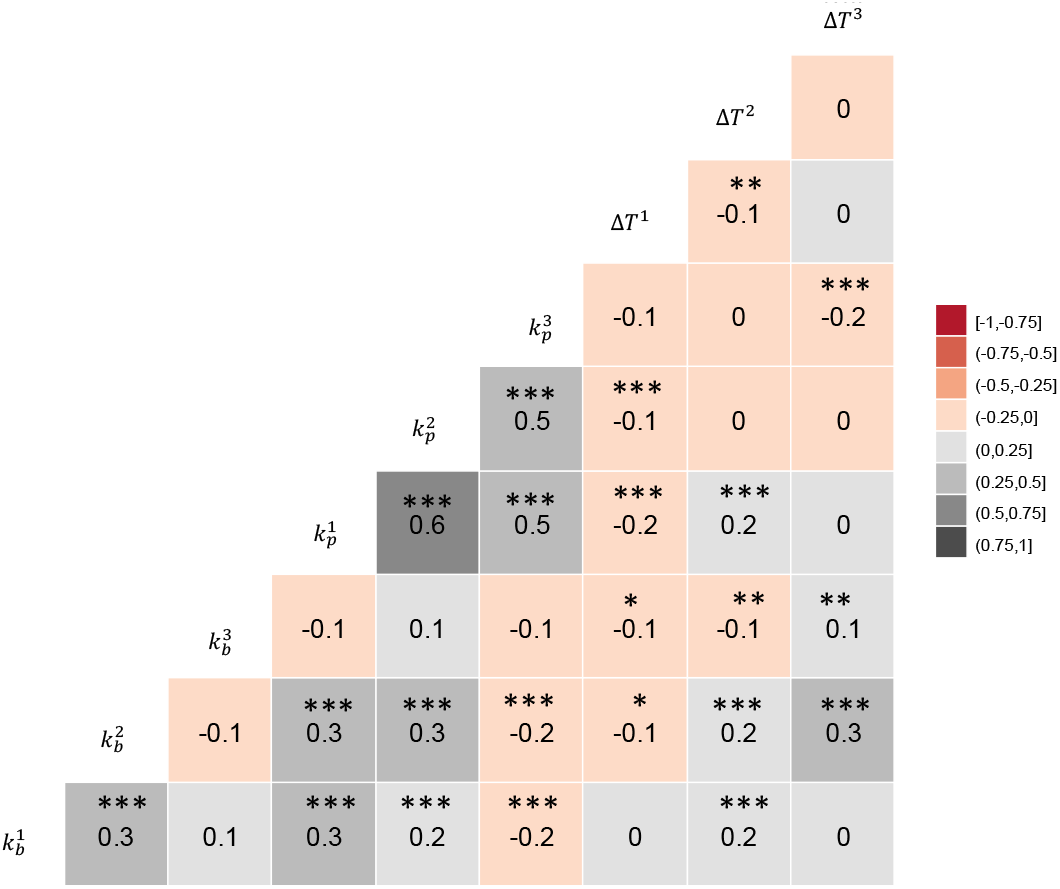
Correlations between variables *k_p_,k_b_* and Δ*T* as estimated by PhenoBR for ewes’ data set used in this study. Grey and red shades stand for positive and negative correlations, respectively. All correlation coefficients and *p*-values are noted. *p*-value, *** < 0.001, ** < 0.01, * <0.05.

Parameters *k_b_, k_p_* and Δ*T* were analysed for the effect of productive cycle, litter size, age at first lambing and the year of BCS measurements (Table 3). Significant effects (*p* < 0.05) of these factors were observed for *k_b_* and *k_p_*. Values of Δ*T* were significantly affected by the productive cycle and the litter size. A decrease was observed for *k_p_* and *k_b_*, as productive cycle increased. Ewes had the longest period of BR mobilization Δ*T* at productive cycle 1 and the shortest Δ*T* at productive cycle 2. As the litter size increased, *k_p_* and Δ*T* increased and *k_b_* decreased. Ewes that lambed for the first time at age one year showed lower *k_b_* and *k_p_* than ewes lambing at age 2 years. The interaction between productive cycle and age of the ewes at first lambing could not be tested owing to uneven distribution of ewes in the classes of each factor. Investigation of the effect of the productive cycle for each class of the age of ewes at first lambing showed similar effects of productive cycle to those described above except for no significant effect of productive cycle on *k_b_* in ewes lambing at age one year (see Supplementary Tables S1 and S2).

**Table 3.** Summary of least-square means for the model variables (standard error) according to the productive cycle, litter size and age of the ewe at first lambing

| | *n* obs | | $k_p$ | $k_b$ | $\Delta t$ |
| --- | --- | --- | --- | --- | --- |
| Productive cycle | 1278 | 1 | 5.14 (0.09) a | 3.25 (0.03) a | 212.96 (1.54) a |
|  | 1204 | 2 | 4.69 (0.09) b | 3.06 (0.03) b | 173.88 (1.52) b |
|  | 521 | 3 | 4.15 (0.10) c | 2.73 (0.05) c | 180.30 (2.32) c |
|  |  | Sign. | *** | *** | *** |
| Litter size | 607 | 1 | 4.23 (0.10) a | 3.25 (0.05) a | 184.09 (2.18) a |
|  | 855 | 2 | 4.64 (0.10) b | 3.04 (0.04) b | 188.21 (1.94) ab |
|  | 830 | 3 | 4.78 (0.10) b | 2.95 (0.04) c | 190.23 (1.86) b |
|  | 712 | 4 | 4.99 (0.10) c | 2.81 (0.04) d | 193.64 (1.97) b |
|  |  | Sign. | *** | *** | * |
| Age at first lambing | 1390 | 1 | 4.45 (0.11) a | 2.88 (0.03) a | 188.04 (1.43) a |
|  | 1614 | 2 | 4.87 (0.10) b | 3.14 (0.03) b | 190.05 (1.35) a |
|  |  | Sign. | *** | ** | NS |
| Year |  | Sign. | *** | NS | NS |
| Productive cycle*litter size |  | Sign. | NS | NS | NS |
| Age at first lambing*litter size |  | Sign. | NS | NS | NS |

*n* obs, number of observations; $k_p$ , rate of body reserve mobilization; $k_b$ , rate of body reserve accretion; $\Delta t$ , duration of body reserve mobilization period; Sign., significance; NS, non-significant. *p*-value, \*\*\* $< 0.001$ , \*\* $< 0.01$ , \* $< 0.05$ . Values of LS means with different letters indicate significant differences between levels of each factor.

A previous study (4), using the ewes involved in the present study, showed that ewes’ BCS variations could be characterized by three classes of trajectory for each productive cycle. Ewes with similar trajectories belonged to the same cluster. Major characteristics of these BCS trajectories including BCS level, BCS loss and gain are given in Table 4 based on previous results reported by Macé et al. (2019). The first productive cycle was characterized by clusters BC1, BC2 and BC3, the second by clusters BC4, BC5 and BC6 and the third by clusters BC7, BC8 and BC9. The two major clusters at production cycles 1 and 2 included 99% and 85% of the ewes, respectively (clusters BC1, 63%; BC2, 36% and BC4, 55%; BC5, 30% respectively). In production cycle 3, the major cluster (BC7) included 76% of ewes (BC8, 13%; BC9,11%).

**Table 4.** Summary of least-square means for the model parameters (standard error) according to clusters of ewes at each productive cycle.

| Cycle | Cluster<sup>1</sup> | BCS level<sup>1</sup> | BCS loss<sup>1</sup> | BCS gain<sup>1</sup> | n ewes | $k_p$ | $k_b$ | $\Delta t$ |
| --- | --- | --- | --- | --- | --- | --- | --- | --- |
| 1 | BC1 | medium | medium | medium | 701 | 5.56 (0.14) a | 3.37 (0.05) a | 212.70 (2.21) a |
|  | BC2 | lowest | lowest | lowest | 432 | 4.99 (0.14) b | 3.12 (0.05) b | 215.83 (2.51) a |
|  | BC3 | highest | highest | highest | 9 | 7.18 (0.64) c | 4.04 (0.37) a | 152.01 (16.72) b |
|  | Sign. |  |  |  |  | *** | *** | *** |
| 2 | BC4 | medium | medium | medium | 593 | 4.82 (0.07) ab | 3.19 (0.04) a | 173.37 (2.22) a |
|  | BC5 | lowest | highest | medium | 328 | 4.98 (0.11) a | 2.83 (0.07) b | 177.04 (3.12) a |
|  | BC6 | highest | lowest | highest | 144 | 4.58 (0.14) b | 3.73 (0.10) c | 170.91 (4.47) a |
|  | Sign. |  |  |  |  | NS | *** | NS |
| 3 | BC7 | medium | highest | medium | 313 | 4.72 (0.10) a | 2.91 (0.07) a | 182.23 (3.30) a |
|  | BC8 | highest | lowest | highest | 48 | 4.15 (0.22) b | 3.72 (0.16) b | 166.15 (7.18) a |
|  | BC9 | lowest | medium | medium | 46 | 5.40 (0.37) a | 2.89 (0.26) a | 186.47 (7.65) a |
|  | Sign. |  |  |  |  | ** | *** | NS |

n ewes, number of ewes in each cluster; $k_p$ , rate of body reserves mobilisation; $k_b$ , rate of body reserves accretion; $\Delta t$ , duration of body reserves mobilization period; Sign., significance; NS, non-significant. $p$ -value, \*\*\* <0.001, \*\* <0.01. Values of LS means with different letters indicate significant differences between clusters within a cycle.
<sup>1</sup> Distribution of ewes in clusters and characteristics of body condition score (BCS) trajectories throughout their productive cycle based on previous results reported by Macé et al. 2018.

Table 4 summarizes the differences in *k_b_, k_p_* and Δ*T* between clusters at each productive cycle. A significant effect (*p* < 0.01) of clusters was observed for *k_p_* at all three parities. For parameter *k_b_*, the effect of clusters was significant at parities 1 and 3. Finally, only the clusters of parity 1 had a significant effect on parameter Δ*T*. The value of *k_p_* was higher for ewes in BC3 at productive cycle 1, which is associated with ewes showing a marked and faster loss of body condition at the beginning of the BR mobilization period, compared to ewes in clusters BC1 and BC2. Values of *k_b_* were higher in ewes belonging to BC1 and BC3 at productive cycle 1, clusters associated with ewes showing higher BR restoring capacity, compared to ewes in BC2. During the second productive cycle, a large difference in *k_b_* was observed between BC5 and BC6. Cluster BC6 was associated with ewes showing the highest BR restoring capacity in the shortest lapse of time, and BC5 was associated with ewes that restored less BR in a longer period. During the third productive cycle, *k_p_* was significantly higher for BC9 and BC7, two clusters with ewes with similar BR dynamics characterized by a marked BR loss. The value of *k_b_* was significantly higher for cluster BC8 corresponding to ewes with the highest BCS levels throughout the third productive cycle.

## Discussion

The aim of the present study was to propose a metric of ewes’ capacity to adapt to their increasing and fluctuating energy requirements through several productive cycles. We exploited the existing database derived from historical and dynamic measurements of BCS in reproductive Romane meat ewes reared in an INRAE experimental farm in France to develop a mathematical model, called PhenoBR. This model converts the individual time series data of BCS into a small number of biologically meaningful synthetic parameters to characterize body reserve dynamics of ewes.

Overall and individual body reserve trajectories had been previously described but without characterizing the individual capacity of ewes for BR mobilization and accretion (3,20). The model developed in the present study characterized each productive cycle *i* with four parameters specific to each ewe: the BR accretion rate 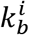, the BR mobilization rate 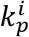, the onset of the mobilization (i.e. onset of the perturbation period) 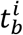 and BR mobilization duration Δ*T^i^*. These parameters, never described before, have the potential to offer new indicators of BR dynamics in meat sheep and potentially in other ruminants. The BR mobilization duration (Δ*T^i^*), estimated in the present study, was consistent with previous results and biological knowledge showing that BR mobilization lasted approximately 180 days in our experimental conditions (i.e., almost 90 days in pregnancy and 90 days in suckling; Macé et al. 2018; 2019).

Although in this study subcutaneous adipose tissue, through values of BCS, was used to illustrate the variations in BRs, it is well known that ewes have other sources of energy stored in their anatomy in the form of adipose tissues (e.g. around internal organs, omental, perirenal, inter- or intra-muscular tissues, etc.). This diversity of adipose tissue sources (locations) can be mobilized in NEB situations (1). Variations in the magnitude of lipid storage in such mostly internal adipose tissues are undetectable by the BCS, which indicates only the status of the subcutaneous fat layer depth. Therefore, when analysing the energy balance status of a given animal, caution must be exercised when interpreting results from BCS alone, as BR mobilization may be in play without a clear change in BCS. However, subcutaneous adipose tissue is considered as the adipose tissue of most interest for investigating BR changes in ruminants since it is reported to be the most labile adipose tissue, and BCS is closely correlated to total body fat content (7). Thus the variables we propose in this study can be considered of high relevance for BR dynamic characterization.

Several models have been developed for converting time-series data into biologically meaningful variables (13,21–26). Some authors have proposed converting the longitudinal data (such as BW, milk yield and BCS) of dairy goats into small number of variables, using a mechanistic model describing priorities of dairy goats throughout their lifespan (11,12). Other models have characterized animal responses to different types of perturbations with approaches mainly based on data (15). The objective of all these models was to detect perturbations that affect animal performance and health, and to characterize the animals’ responses during such periods. When data were available with high frequency, the models were able to detect periods of unknown perturbations. The originality of PhenoBR compared to the existing models is its use of a combination of a data-driven model (via FDA) and a dynamic model to detect perturbation periods despite the limited number of available BCS records, and to subsequently characterize ewe’s response. Using the data analysis approach by fda helped to decrease the number of variables to be estimated for the dynamic model and thereby increase the robustness of the parameter estimation process (convergence of the process for all animals).

Several biological factors had a significant effect on 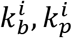 and Δ*T^i^* and were similar to those affecting BCS when considered at a single time-point (20,27–29).

The decrease observed for BR mobilization rate *k_p_* and BR accretion rate *k_b_* (Table 3) as the parity increased, indicates that body condition losses were more marked at the first productive cycle during the BR mobilization period and gained more during the BR accretion period, compared to productive cycles 2 and 3. However, results showed that for ewes lambing at age one year (Supplementary Table S1), the effect of parity on their recovery capacity *k_b_* was not significant. Given that they decrease their *k_p_* as their parity increased, this suggests better performance in ewes lambing at an earlier age.

According to BR mobilization and accretion rates, younger ewes at first lambing lost less body condition and recovered at a lower rate than older ewes. This may be linked to the fact that ewes at age one year are still growing and allocate less energy to adipose tissues to continue their growth and assure their next reproduction cycle. The “round” nature of such growth in younger ewes could induce biases in the BCS assessment as during palpation of the dorsal region, the operator may recognize a larger and confounded mass of muscle and adipose tissues. This is likely the case for the first BCS point in one part of our ewes present in the database (Supplementary Table S2) for which the first BCS could be overestimated in comparison with the same BCS or energy balance status in the following parities. The slope between this first BCS value before mating and the following may be biased, which could affect *k_p_* results in our model (i.e. most drastic BCS variation).

Our results are consistent with those reported by Macé et al. (3) showing similar effects of parity and age at first lambing when considering BCS changes between successive physiological stages. The BR mobilization and accretion rates estimated in the present study indicated that ewes were able to mobilize more BR as litter size increased, while the BR accretion was less marked for ewes with larger litter size. The increase in BR mobilization rate was expected because of the classical related higher energy requirements for ewes suckling multiple litters. Similar effects of litter size on BR losses during pregnancy and suckling had been previously reported when considering BCS differences between some physiological stages (3,20,28,30). However, the decrease in BR accretion rate with the increase in litter size conflicted with previous results showing higher BR accretion in ewes with higher litter size (3). This discrepancy may be related to the fact that in the modelling, BR accretion starts from the beginning of the perturbation, i.e. BR accretion rate was not only considered in the recovery period but also during the theoretical perturbation period in which short anabolic and catabolic reactions could be encompassed.

Interesting positive correlations were found for BR mobilization rates (*k_p_*) between parities, suggesting that ewes maintained their biological capacity for BR mobilization across productive cycles. However, correlations between parities for the other two parameters 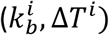 suggested that BR accretion rate and BR mobilization duration varied across productive cycles. This could be due to higher environmental effects of each cycle (i.e. feed availability, number of lambs previously suckled, etc.) on BR accretion rate and BR mobilization time. Correlations were also found between BR mobilization rate and BR accretion rate suggesting both processes are biologically linked as previously claimed by Chilliard et al. (1). This result was consistent with high genetic correlations found previously between BR mobilization and accretion in the same dataset, which indicated that ewes exhibiting high level of BR mobilization also exhibited high level of BR accretion (Macé et al., 2018).

The variability in BCS profiles throughout ewes’ lifespan had already been investigated by the presence of clusters in each productive cycle (4). Each of these clusters were characterized by BR trajectories differing in the level of BR and/or shape of BR changes through the productive cycle. In the present results, we found consistency between average values of BR mobilization and accretion rates and characteristics of BR profiles in clusters. Differences between clusters found in the present study for BR accretion rate *k_b_* and BR mobilization rate *k_p_* suggest that such parameters could be used for ranking animals according to their BR dynamics. Thus PhenoBR offers an opportunity to investigate ewes’ variability of response at the individual level.

## Conclusions

The existing inter-individual variability for the parameters quantified with PhenoBR, BR mobilization and accretion rates and duration of BR mobilization make them potentially useful for classifying individuals according to their BR dynamics. Such parameters could be used by farmers for animal management, for instance by adapting feeding systems according to the individual characteristics of animals in terms of BR dynamics and the seasonality of feed and forage availability. Tthese parameters could also be used by geneticists as criteria to refine and reinforce future animal breeding programs, including selection strategies for more robust and resilient animals. Combining these new traits with other related metabolites and metabolic hormone data, known to be related to BCS and therefore to the BR mobilization and accretion processes, will take us further in our understanding of mechanisms underlying animal response to NEB perturbations. Altogether, in the perspective of working towards resilient farming systems, PhenoBR will support our approach for taking advantage from ruminants’ adaptive capacities when facing successive NEB periods related to exigent physiological events or other environmental perturbations affecting nutrient availability and nutritional balances.

## Acknowledgements

The authors are indebted to all the staff of ‘La Fage’ experimental farm for managing the experimental flock as well as for their active role in collecting experimental data. The development of the web interface benefitted from the interactions provided by the INRAE PHASE project PhenoBR.

## Availability of data and material

The R script of ODE model and the dataset used for this study are available in public repository Zenodo (http://doi.org/10.5281/zenodo.4300412). The software is freely available on http://adaptive-capacity.herokuapp.com/.

## Contributions

E.G. and D.H. designed the study and contributed to produce the database used in the current work. M.T. and T.M. developed the model and data analysis procedures. G.K. and M.T. developed the web interface. M.T drafted the manuscript. All authors contributed to the analysis and interpretation of the results. All authors read and approved the final manuscript.

