## Supplementary materials for "PhenoBR: a model to phenotype body condition dynamics in meat sheep"

Tiphaine Macé[1], Eliel Gonzalez Garcia[2], György Kövér[3], Dominique Hazard[1] and Masoomeh Taghipoor[4*]

[1] GenPhySE, Université de Toulouse, INRAE, ENVT, F-31326 Castanet-Tolosan, France
[2]SELMET, INRAE CIRAD, Montpellier SupAgro, Univ Montpellier, 34060 Montpellier, France
[3] Szent István University, Kaposvár Campus H-7401 Kaposvár, Guba S. u. 40 , Hungary
[4]Université Paris-Saclay, INRAE, AgroParisTech, UMR Modélisation Systémique Appliquée aux Ruminants, 75005, Paris, France

**Supplementary hypothesis S1**

For productive cycle 1, if $t_b$ is not included between days 0 and 160 (i.e. approximatively between mating and lambing days) and $t_e$ is not included between days 160 and 330 (i.e. approximatively between lambing and post-weaning):

- If no BCS maximum found, then $t_b$=0;
- If no BCS minimum were found between days 160 and 330 but there is a BCS minimum before day 330 then $t_e$=day where BCS is minimal;
- If no BCS minimum found, then $t_e$=360;
- If two minimums found during the perturbation time, then $t_e$= day where BCS is minimal.

For productive cycle 2, if $t_b$ is not included between days 360 and 520 (i.e. approximatively between mating and lambing days) and $t_e$ is not included between days 520 and 690 (i.e. approximatively between lambing and post-weaning):

- If no BCS maximum found between days 360 and 520 but there is a BCS maximum after day 520 then $t_b$=day where BCS is maximal;
- If no BCS minimum were found between days 520 and 690 but there is a BCS minimum before day 520 then $t_e$= day where BCS is minimal;
- If no BCS maximum found, then $t_b$=360;
- If no BCS minimum found, then $t_e$=720;
- If two BCS maximums found before the perturbation time, then $t_b$ = day where BCS is maximal;
- If two minimums found during the perturbation time, then $t_e$== day where BCS is minimal.

For productive cycle 3, if $t_b$ is not included between days 720 and 880 (i.e. approximatively between mating and lambing days) and $t_e$ is not included between days 880 and 1050 (i.e. approximatively between lambing and post-weaning):

- If no BCS maximum found between days 720 and 880 but there is a BCS maximum before day 720 or after day 880 then $t_b$= day where BCS is maximal;
- If no BCS minimum found between days 880 and 1050 but there is a BCS minimum before day 880 then $t_e$= day where BCS is minimal;
- If no BCS minimum found, then $t_e$=1050;
- If no BCS maximum found, then $t_b$=720.

Table S1. Summary of least-squares means for the model parameters (standard error) according to the productive cycle and significance of litter size, year and productive cycle x litter size interaction effects in ewes performing first lambing at 1 year old.

| | | $k_p$ | $k_b$ | Perturbation length ($\Delta t$) |
|---|---|---|---|---|
| n obs | | 1390 | 1390 | 1390 |
| productive cycle | 1 | 4.53 (0.17) a | 2.92 (0.15) a | 205.39 (7.89) a |
| | 2 | 4.49 (0.06) a | 2.96 (0.05) a | 166.21 (2.40) b |
| | 3 | 3.80 (0.09) b | 2.96 (0.07) a | 165.87 (3.89) b |
| | Sign. | *** | NS | *** |
| litter size | Sign. | *** | *** | NS |
| year | Sign. | *** | *** | *** |
| productive cycle*litter size | Sign. | NS | NS (0.07) | ** |

n obs, Number of observations; $k_p$, Rate of body reserves mobilization; $k_b$, Rate of body reserves accretion; $\Delta t$, duration of body reserves mobilization period; Sign., Significance; NS, non-significant. Pvalue, *** < 0.001, ** < 0.01, *<0.05 ; Values of LSmeans with different letters indicate significant differences between levels of each factor.

Table S2. Summary of least-squares means for the model parameters (standard error) according to the productive cycle and significance of litter size, year and productive cycle x litter size interaction effects in ewes performing first lambing at 2 years old.

| | | $k_p$ | $k_b$ | Perturbation length ($\Delta t$) |
|---|---|---|---|---|
| n obs | | 1614 | 1614 | 1614 |
| productive cycle | 1 | 5.37 (0.11) a | 3.48 (0.08) a | 204.15 (3.76) a |
| | 2 | 4.79 (0.12) b | 3.15 (0.08) b | 177.65 (3.96) b |
| | 3 | 4.33 (0.13) c | 2.69 (0.09) c | 171.00 (4.65) b |
| | Sign. | *** | *** | *** |
| litter size | Sign. | *** | *** | *** |
| year | Sign. | *** | *** | *** |
| productive cycle*litter size | Sign. | NS | NS | NS |

n obs, Number of observations; $k_p$, Rate of body reserves mobilization; $k_b$, Rate of body reserves accretion; $\Delta t$, duration of body reserves mobilization period; Sign., Significance; NS, non-significant. Pvalue, *** < 0.001, ** < 0.01, *<0.05 ; Values of LSmeans with different letters indicate significant differences between levels of each factor.

Table S3. Summary of least-squares means for the model parameters (standard error) according to the clusters of adjusted BCS

| | | n obs | $k_p$ | $k_b$ | Perturbation length |
|---|---|---|---|---|---|
| Productive cycle 1 | BC10 | 1037 | 5.16 (0.13) a | 3.33 (0.04) a | 213.74 (1.85) a |
| | BC11 | 105 | 6.36 (0.20) b | 2.90 (0.10) b | 214.29 (4.94) a |
| | Sign. | | *** | *** | NS |
| n obs | | | 1069 | 1069 | 1069 |
| Productive cycle 2 | BC12 | 822 | 4.80 (0.06) a | 3.15 (0.04) a | 176.65 (2.02) a |
| | BC13 | 247 | 4.96 (0.11) a | 3.29 (0.08) a | 166.45 (3.50) b |
| | Sign. | | NS | NS | * |
| n obs | | | 414 | 414 | 414 |
| Productive cycle 3 | BC14 | 235 | 4.68 (0.12) a | 2.98 (0.08) a | 185.42 (3.71) a |
| | BC15 | 179 | 4.57 (0.12) a | 3.02 (0.08) a | 175.30 (3.79) b |
| | Sign. | | NS | NS | * |

n obs, Number of observations; Pvalue, *** < 0.001, *<0.05; Sign., Significance; NS, non-significant.

Productive cycle 1:
- BC10: lowest BCS loss and BCS gain;
- BC11: highest BCS loss and BCS gain.

Productive cycle 2:
- BC12: lowest BCS loss and intermediate BCS gain;
- BC13: highest BCS loss and intermediate BCS gain.

Productive cycle 3:
- BC14: intermediate BCS loss and BCS gain;

BC15: intermediate BCS loss and BCS gain

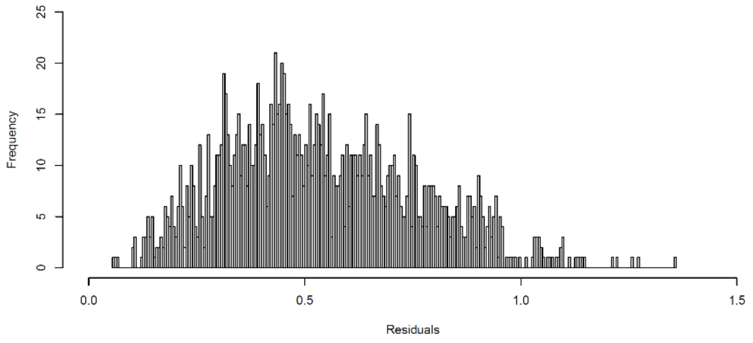


Figure S1. Histogram of the objective function ($f_{obj}$). Only a limited number of ewes with $f_{obj}$>1 could be observed.
